# A Multidimensional Arithmetical Ontology for High-Parameter Cytometry: Gaussian Primes and Cryptographic Hashing for Cell Population Standardization

**DOI:** 10.64898/2026.10.14.758936

**Authors:** Alice Yue, Cecile Masquelier, Antonio Cosma

**Affiliations:** Liangzhu Laboratory, Zhejiang University School of Medicine, Zhejiang, China; National Cytometry Platform, Translational Medicine Operations Hub, Luxembourg Institute of Health, Esch-sur-Alzette, Luxembourg

**Keywords:** flow cytometry, cell population standardization, Gaussian primes, cryptographic hashing, ontology, SHA-256, gating strategy, Prime Population System, UniProt, machine learning interoperability

## Abstract

**Background:** The rapid expansion of high-dimensional single-cell technologies has outpaced the development of standardized metadata infrastructure. Current text-based ontology for cell populations and gating strategies lacks computational rigor, hindering process automation and cross-study reproducibility. Our previous frameworks, the Prime Population System (PPS) and the Unique Gating Identifier (UGI), proposed arithmetical solutions but were limited by biological fragility (no inherent link between a protein and its structural isoforms) and by computational bottlenecks caused by exponential integer growth.

**Objective:** To develop a scalable, multidimensional arithmetical ontology that unambiguously identifies cell populations and hierarchical gating strategies.

**Methods:** We transition the marker dictionary from the one-dimensional real axis (ℝ) to the complex plane of Gaussian integers (*a*+*bi*). The reviewed human proteome of the UniProt Knowledgebase is paired with Gaussian primes satisfying *a* > *b* > 0. Each protein receives a contiguous reserved block of 64 primes: slot 0 addresses the protein with isoform unresolved, and slot *k* addresses UniProt isoform *k*, the slot index being read directly off the accession. Carbohydrate epitopes, which have no UniProt entry, are paired through the CD nomenclature instead: in a CD region beyond the 80,000 blocks reserved for the proteome, each CD number receives a block of 1,024 primes, with the slot read off the suffix of the designation. A bipartite encoding represents positive (a+bi) and negative (b+ai) expression states through Conjugate Pair Symmetry. To avoid the integer explosion of Gödel exponentiation, hierarchical gating sequences are held as vectorized arrays of complex numbers and compressed by SHA-256 deterministic hashing.

**Results:** Pairing against the human proteome assigns 53,296 identifiers, 20,431 protein-level and 32,865 isoform-level. Every isoform description resolves under a single rule, including the 329 descriptions whose accession prefix differs from the entry that cites them. The 19 carbohydrate designations of the CD list, CD57 among them, receive identifiers in the CD region. Localized vectorization with SHA-256 hashing yields immutable 64-character hexadecimal fingerprints for entire gating strategies, resolving the integer explosion problem.

**Conclusion:** By combining Gaussian primes with hashing, Gaussian Prime Cytometry provides a scalable, computationally lightweight and biologically accurate identifier system. The system eliminates nomenclature ambiguity and facilitates machine learning applications and centralized global cytometry databases.

## 1. Introduction

### 1.1 Background and the standardization challenge

The continuous evolution of single-cell technologies allows the simultaneous measurement of dozens to hundreds of cellular parameters (i.e. markers), yet the field suffers from a critical lack of ontological standardization. Experiment metadata, such as gating strategies and cell types, are described using subjective, text-based nomenclature. This ambiguous nomenclature makes it exceedingly difficult to share, reproduce or computationally query metadata across software platforms, data repositories and publications.

### 1.2 Foundational mathematical frameworks

#### Cell population annotation — the Prime Population System (PPS)

To eliminate the errors inherent in traditional naming conventions, Cosma (1) introduced the Prime Population System, a method rooted in the fundamental theorem of arithmetic. By assigning a unique natural prime *P* to each biological marker, any multiplexed cell population is unambiguously identified by the product of its expressed *n* marker primes (*p*_*i*_). That product is unique and can be unambiguously factorized to recover the original marker prime set.

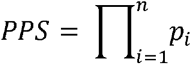

#### Gating strategy annotation — the Unique Gating Identifier (UGI)

Building on the PPS, Cosma (2) extended the framework to encode the hierarchical, sequential nature of gating strategies using Gödel numbering. Each sequential gate is assigned a consecutive prime (*P*_j_) raised to the power of its PPS number; the product of these terms constitutes the UGI, a unique arithmetic fingerprint for any existing or future gating strategy with *m* sequential gates:

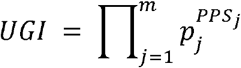

### 1.3 Current limitations of biological fragility and computational bottlenecks

While mathematically rigorous, scaling this model into a globally applicable software standard presents three distinct challenges.

#### Dictionary inflation

Mapping the entire human proteome with standard natural primes produces rapidly escalating base numbers, which makes calculations hard to handle even for small marker panels.

#### Biological fragility

Linear prime assignment fails to capture the relationship between a parent molecule and its structurally related isoforms (for example CD45, CD45RA, CD45RO). Furthermore, while negative expression was encoded by squaring the marker’s prime (*p*^*2*^), this convention creates no intrinsic mathematical link between the positive (*p*^*1*^) and negative (*p*^*2*^) states: the relationship is a convention, not a property of the number itself.

#### Integer explosion

Because Gödel numbering raises each gate’s prime to the power of its PPS number, a single gate on ten markers already yields an integer of more than a billion digits.

### 1.4 Proposed extensions

This manuscript proposes a unified, scalable architecture that addresses these limitations. By introducing Gaussian integers (3), bipartite state symmetry, reserved isoform blocks and SHA-256 cryptographic hashing, we present Gaussian Prime Cytometry, a scalable and biologically accurate identifier system.

## 2. Methods

### 2.1 Construction of the Gaussian prime dictionary

The prime dictionary is a pairing of two lists, each totally ordered by a key that already exists in its own source. Because each list is ordered independently and objectively, pairing them by position requires no curation, no expert consensus and no registry: the *n*-th reviewed human protein is paired with the *n*-th block of 64 admissible Gaussian primes by construction (Figure 1A). The rules that follow specify the two orderings, the pairing itself, and the resolution of every case that arises in the source data.

**Figure 1.**
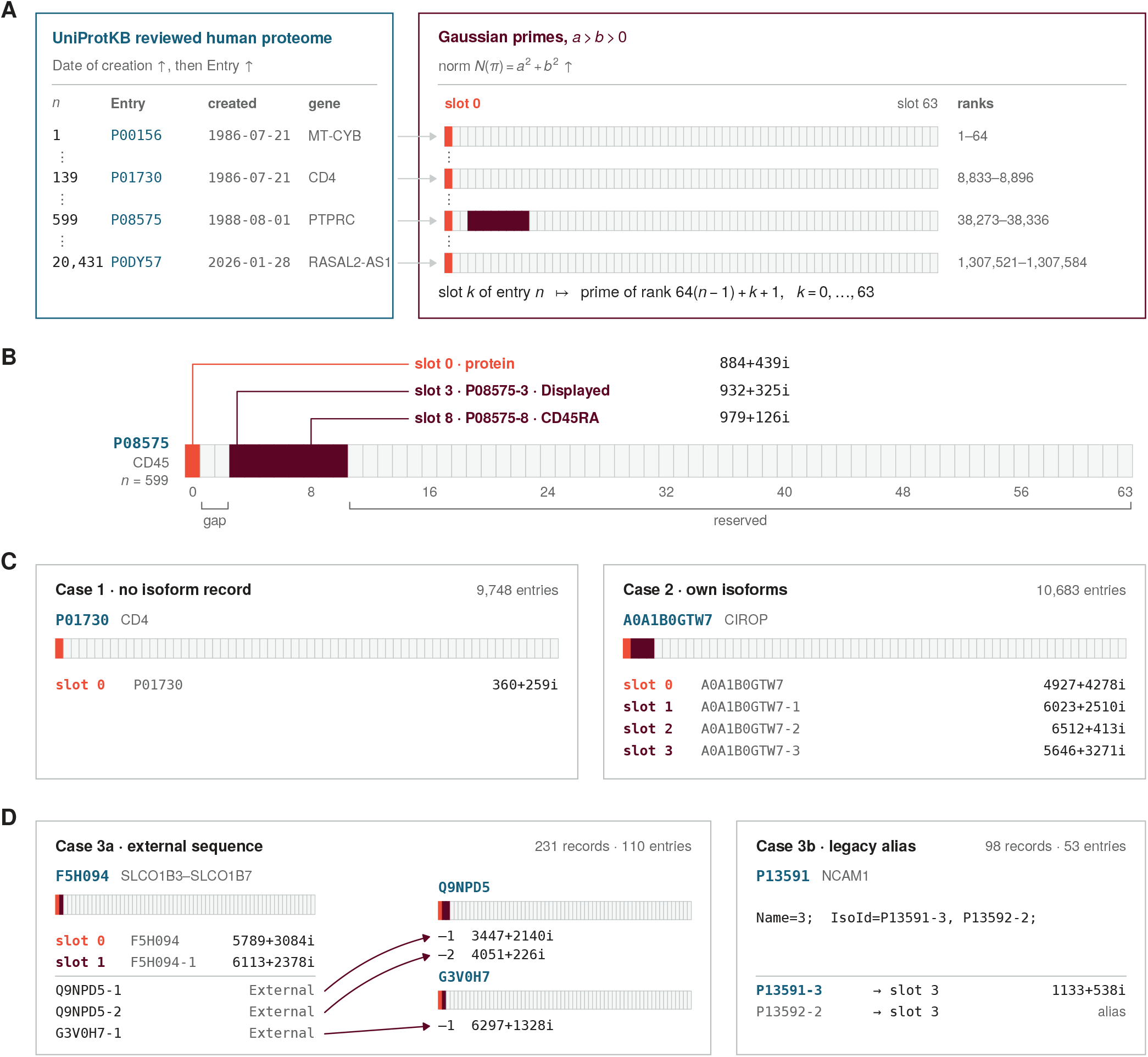
The pairing rules of the Gaussian prime dictionary. **(A)** The dictionary pairs two independently ordered lists. The reviewed human proteome of UniProtKB is ordered by date of creation ascending, then by primary accession ascending under plain byte comparison; Gaussian primes satisfying a > b > 0 are ordered by ascending norm. Each entry owns a contiguous block of 64 primes, so slot *k* of entry *n* is the prime of rank 64(*n*−1)+*k*+1. **(B)** Anatomy of one block, shown for CD45 (P08575, entry 599). Slot 0 addresses the protein with isoform unresolved; slot *k* addresses UniProt isoform *k*, the index being read directly off the accession. CD45 has no isoform −1 or −2 and its Displayed sequence sits at −3, illustrating why slot 1 cannot serve as the protein-level address. Gaps are retained and never reused. **(C)** The two ordinary cases, in which every IsoId prefix is the entry itself. **(D)** The special case in which an IsoId prefix differs from the entry. External sequences resolve to the owner’s block and mint no new prime; legacy aliases are synonyms resolved by a shared isoform description rather than by matching suffix. Counts are for the reference release (retrieved 23 March 2026, 20,431 entries).

#### 2.1.1 The reference dictionary and its ordering

The reference dictionary is the reviewed (Swiss-Prot) section of the UniProt Knowledgebase (4) restricted to *Homo sapiens*, retrieved on 23 March 2026 and comprising 20,431 entries. This snapshot is designated the **reference release;** all identifiers reported here are allocated against it.

Entries are ordered by two keys applied in sequence:

1. Date of creation, ascending — the date on which the accession first entered the knowledgebase;
2. Entry, the primary accession, ascending — compared as a plain lexicographic byte sequence.

The first key is chosen because it is immutable and monotone. A protein’s creation date never changes, and an entry admitted after the reference release necessarily carries a later date. New entries therefore append to the end of the list rather than being interleaved, and no existing index is displaced. The second key resolves entries sharing a creation date — of which there are many — in a manner likewise fixed for all time.

The second key is a plain string comparison, **not** a numerically aware “natural” sort. Implementations must specify plain byte comparison explicitly.

The ordering is total and deterministic. Both keys are present for every entry, and the pair (creation date, accession) is unique, so no further tie-break is ever required. The reference release spans 379 distinct creation dates. An entry’s position in this ordering is its **index**, written *n*; the word **rank** is reserved for positions in the prime list (§2.1.2).

##### Index allocation policy

An index is allocated on first observation. An accession receives the next free index in the release in which it first appears in the reviewed human set, and that index is permanent. An accession that subsequently leaves the reviewed set — by merging, demotion or deletion — retains its index as a tombstone; the index is never reused and never reassigned. This policy is what makes the dictionary append-only, and it closes the one mechanism by which a returning accession could otherwise disturb existing indices. Because an index is allocated when an accession is first observed, rather than recomputed from the creation date at each release, an accession that re-enters the reviewed set carrying its original creation date cannot displace indices already allocated.

#### 2.1.2 The Gaussian prime list and its ordering

The fundamental theorem of arithmetic extends to Gaussian integers ℤ[*i* ]every nonzero Gaussian integer that is not a unit factors uniquely into Gaussian primes, up to the order of the primes and multiplication by units (5, Theorem 215). Two Gaussian integers are associates if they differ only by multiplication by a unit factor (1,−1,*i*,−*i*).

The prime list must therefore be restricted so that no two admitted primes are associates, and so that no admitted prime is an associate of the mirror image used to encode negative expression (§2.2.1).

A Gaussian prime π*= a + bi;* is admitted to the prime list *D* if and only if

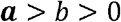

This constraint means we select exactly one representative from each set of associates in the first quadrant, strictly below the line *y= x*. This also means that we exclude inert primes (b = 0) and the prime 1 + i, of the form a = b. The negative counterpart of a marker is obtained by swapping the real and imaginary parts (§2.2.1), so the counterpart of an inert prime *a +* 0*i;*, with *a* ≡*3* (*mod 4*), would be *0+ ai*. But *0 + ai = i ·* (*a + 0i*),so the two are associates: they differ only by the unit *i*. Because factorization over ℤ[*i* ] is unique only up to associates, the positive and negative states of an inert marker would be indistinguishable under factorization. Admitting inert primes would break Conjugate Pair Symmetry. The mirror image of a prime where a = b will be indistinguishable from itself when the real and imaginary part are swapped and it will not be possible to use them to encode negative markers.

The Gaussian primes in *D* are organized in ascending order of their norm *N(*π*) = a*^*2*^*+ b*^*2*^. Since each admissible norm is a distinct rational prime, the norms are pairwise distinct, and the ordering is total without any tie-break — mirroring the property that makes the proteome (all possible markers) ordering deterministic. A prime position in this ordering is its **rank**.

A list of the first 1,920,000 admissible primes was generated for this work, running from 2 *+* 1*i;* (norm 5) to 8060 *+* 199*i* (norm 65,003,201), with no component exceeding 8,062. The list was validated in four independent ways: every element satisfies *a> b > 0* and *N(*π*)1* ≡ (*mod 4*); norms are strictly increasing and unique; an independent primality test on a random sample of 3,000 norms returned no failures; and the list agrees exactly, in both membership and order, with an independently generated enumeration of Gaussian primes to norm 10^6^ over the shared range (39,175 elements).

#### 2.1.3 The pairing rule

One constant is fixed at publication: the block depth ***B=* 64**. The ordered prime list is partitioned into contiguous blocks of *s* primes, and the entry of index *n* owns the block spanning ranks B · (*n*−1)+1 through B · *n* (Figure 1A-B).

Within a block, slot *k* of entry *n* is paired with the Gaussian prime of rank

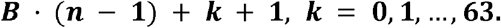

The slot index is read directly off the UniProt identifier: *k* is the integer following the hyphen, or 0 when there is none. Thus, P08575 addresses slot 0 and P08575-3 addresses slot 3 (Figure 1B). Nothing — no lookup, no interpretation, no annotation judgement — intervenes between the identifier and its slot. The highest isoform suffix anywhere in the reference release is 37, carried by CACNA1C (Q13936), so B = 64 leaves 26 numbered slots spare even for the deepest entry.

#### 2.1.4 The protein-level address (slot 0)

Slot 0 addresses the protein itself, with isoform unresolved. It is the identifier used when an antibody clone is pan-reactive, and equally when its isoform reactivity has not been determined. The two are deliberately not distinguished: whether a clone is genuinely pan-reactive or merely uncharacterized is a property of the reagent, not of the protein, and encoding that distinction in the identifier would force an annotation judgement at the moment of encoding — precisely the subjectivity the framework exists to remove. It belongs to the antibody metadata.

Reserving slot 0 rather than overloading slot 1 is forced by the data. In the reference release, **477 entries** carry their Displayed (representative) sequence at a suffix other than 1, in one case as high as 15, and **116 entries** possess no isoform −1 at all, the suffix having been retired. For those entries slot 1 holds either a non-representative variant or nothing. CD45 (P08575) exhibits both pathologies: its lowest suffix is 3 and its Displayed sequence sits there (Figure 1B). Under a slot-1 convention, the protein most central to cytometric gating would have had no protein-level identifier at all.

Separation is also semantically necessary. “This protein” and “this specific splice variant” are different claims; collapsing them into a single identifier is exactly the homonymy this framework is designed to eliminate.

#### 2.1.5 Resolution of isoform descriptions

The UniProt ALTERNATIVE PRODUCTS field opens with a header giving the event type and the number of named isoforms, followed by one isoform description per isoform: a run of semicolon-separated key–value pairs that begins with Name=, ends with `Sequence=` and lists one or more `IsoId`s. Each `IsoId` becomes one database record, placed by the following rule.

Write each IsoId as *P*-*k*, where *P* is the accession before the hyphen and *k* is the integer after it, and let *E* be the entry where isoform descriptions are listed. The `IsoId` is placed by comparing *P* with *E*:

**(Own IsoId)** If *P* = *E*, the `IsoId` is one of *E*’s own isoforms. It occupies slot *k* of *E*’s block and takes the Gaussian prime of that slot. Only own `IsoId`s occupy slots, so a block holds isoforms of its own entry and of no other. A suffix is used only once within an entry, so no slot is occupied twice. For example, `P08575-3`, listed by `P08575`, occupies slot `3` of the `P08575` block.

**(Foreign IsoId)** If *P* ≠ *E*, the `IsoId` carries the *P* of another accession: *E* lists it, but the isoform is not numbered within E. It occupies no slot in *E*’s block, and its suffix *k* says nothing about *E*’s slots: it is a position in *P*’s numbering. What the `IsoId` refers to depends on whether *P* is a current entry of the reference set.

**(Borrowed)** If *P* is a current entry, the `IsoId` addresses slot *k* of P’s block without occupying it. *E* borrows *P*’s identifier, and no new prime is created. For example, `Q9NPD5-1`, listed by `F5H094`, addresses slot 1 of the `Q9NPD5` block; slot 1 of the `F5H094` block belongs to a different isoform, `F5H094-1`.

**(Retired)** If *P* has been retired and is no longer an entry of the reference set, it owns no block, and the `IsoId` addresses none. For example, `P13592-2` appears in an isoform description of `P13591` together with `P13591-3` and is read as `P13591-3`. `P13592-2` is just another alias of `P13591-3`, it doesn’t get its own prime.

Applied to the reference release, these rules classify every IsoId without exception. The rules are stated for a single IsoId, but identifiers are looked up by entry, so we also describe the entries themselves. Grouped by the kind of IsoIds they list, the 20,431 entries fall into three cases (Figure 1C, 1D). The cases show what the rules do to an entry’s block. Because their counts add up to every entry and every IsoId in the release, they confirm that no IsoId escapes the rule.

##### Case 1

**has no isoform description** (9,748 entries, 47.7%). The entry has no alternative entries or isoforms. Slot 0 is the entire identifier and slots 1–63 remain reserved. An example of this is CD4 (`P01730`).

##### Case 2

**has isoform description and *P*** = ***E*** (10,527 entries, 51.5%; 34 with a single isoform, 10,493 with two or more). Every alias prefix is the entry itself, and slot *k* carries isoform −*k*. Slot indices need not be contiguous, and a gap records a retired isoform that is never reused.

##### Case 3

**has an alias whose prefix is not the entry and *P* ≠ *E*** (156 entries, 0.8%). Two sub-cases occur, distinguished by whether the isoform description also holds an IsoId with the entry’s own prefix.

*Case 3a — external sequences* (231 descriptions across 110 entries, holding 247 `IsoIds`), flagged Sequence=External. None of these descriptions’ `IsoId`s contain their entry’s accession (all *P* ≠ *E*): the sequence is defined in another entry and retains that entry’s accession, as is typical of read-through and fusion transcripts that reuse component-gene isoforms. All its IsoIds are foreign. Of the 247 `IsoIds`, 232 carry a current accession and address a slot their owner already allocates, under the identical `IsoId` string, so there are no collisions and no additional primes are minted; the other 15 carry a retired accession and address no block. Every one of the 231 descriptions therefore reaches an identifier its owner holds. `F5H094`, the `SLCO1B3`–`SLCO1B7` read- through protein, illustrates this: its three external descriptions resolve into the blocks of `Q9NPD5` and `G3V0H7` (Figure 1D).

*Case 3b — legacy aliases* (98 descriptions across 53 entries, each holding the entry’s own `IsoId` and one alias). Each description has one own IsoId (*P* = *E*) plus one alias with a different prefix. 96 descriptions use retired accessions left behind after UniProt merged entries, because `IsoId` codes stay stable when an accession is replaced. Their own `IsoId` occupies the entry’s slot; the retired alias addresses no block and is a synonym of the own identifier through the shared description. The two aliases do not necessarily share the same suffix *k* (e.g., `P85298-1` is aliased by `Q9NSG0-6`, and `P13591-3` by `P13592-2`), so a legacy alias is resolved by sharing a description, never by matching suffix number.

One sequence holds two identifiers. The chimeric `ERCC6–PGBD3` protein carries two `IsoIds` whose accessions are both current entries, `P0DP91-1` and `Q03468-2`, and each entry allocates its own; these are the remaining two Case 3b descriptions. Where `PGBD3` (`Q8N328`) cites the chimera as an external sequence, its two `IsoIds` lead to the same two identifiers. No other sequence in the release carries current `IsoIds` in two entries.

The rule rests on one principle: **the accession prefix *P*, not the citing entry, determines which block an identifier inhabits**. A foreign suffix is an index within the owner’s numbering and carries no meaning in the citing entry’s block.

### 2.2 Resolution of expression states

#### 2.2.1 Bipartite encoding of expression state

Gaussian primes make it simple to record the positive and negative versions of a marker. By default, the Gaussian prime assigned to each marker represents the marker’s positive state (e.g. CD45+). Given a Gaussian prime *a+ b;* where *a> b > 0*, we get the marker’s negative counterpart (e.g. CD45-) by swapping the real and imaginary components *a* and *b*. We call this the principle of **Conjugate Pair Symmetry** (Figure 2). Because the prime list is restricted to *a> b > 0* (§2.1.2), the positive and negative forms of every marker are distinct and non-associate, and both are recoverable by factorization over ℤ[*i*]. When a cell population is defined, its identifier is the product of all expressed complex numbers. This yields a unique composite Gaussian integer for any cell type, completely factorizable into its base markers, states and isoforms.

**Figure 2.**
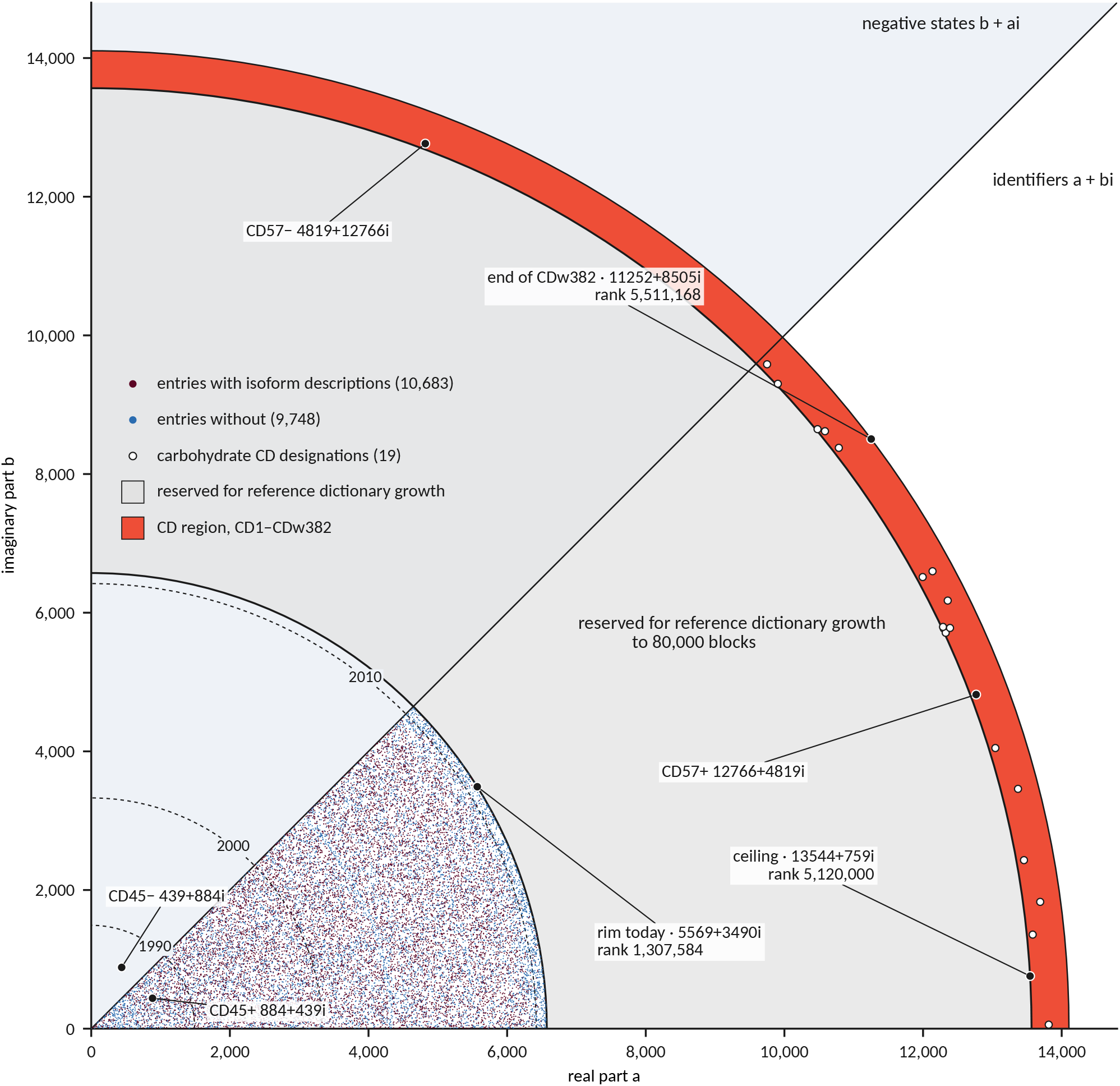
The reference release on the complex plane. Each point is the protein-level (slot 0) Gaussian prime of one of the 20,431 entries of the reviewed human proteome; red, entries with isoform descriptions (10,683); blue, entries without (9,748). All identifiers lie in the region a > b > 0; the region b > a > 0 holds the corresponding negative states b + ai, shown for CD45 (884+439i and its mirror 439+884i). Because entries are ranked by creation date and primes by norm, radius increases with creation date: dashed arcs mark the outermost entry created by the end of each year shown. The grey region is reserved for the growth of the reference dictionary, up to 80,000 blocks (§2.4). The orange region is reserved for CD nomenclature. The 19 carbohydrate identifiers are in white. CD57 (12766+4819i) and its mirror (4819+12766i) are shown in black.

#### 2.2.2 One quadrant exclusion encoding

A quadrant gate on a plot of two markers selects one combination of their states, and its identifier is the product of the two marker states (§2.2.1). Some populations, however, pool the cells of three quadrants and are defined by the one they lack. Non-naive T cells can be gated as every T cell outside the CCR7+ CD45RA+ quadrant of naive cells (6), pooling central memory, effector memory and TEMRA cells, and monocytes as the HLA- DR+ cells outside the CD14− CD16− quadrant, pooling the three monocyte subsets (7). Such a gate is drawn as an L-shaped polygon or obtained with the NOT operation of the analysis software; we call it an **L-gate**. No product of marker states can encode it, because a product lists the states its cells share, and the three quadrants share none.

We encode an L-gate as −**1 times the identifier of the quadrant it excludes**. Multiplying by the unit −1, a half-turn about the origin, changes none of the prime factors, so the excluded markers keep their states and the −1 alone records their exclusion. The L-gate of non-naive T cells is thus −273056461926722−1797474067315335i.

The −1 survives factorization. By unique factorization (§2.1.2), every nonzero Gaussian integer is a unit times a product of primes taken in the first quadrant of the complex plane, and both the unit and the primes are unique. Dictionary primes and their mirrors all lie in that quadrant, so the identifiers of marker states, quadrants and populations carry the unit 1, and L-gates carry −1. Dividing an L-gate by its primes returns the excluded markers in their states, with −1 as the remainder, so each L-gate factorizes back into the single gate it labels, never equals the identifier of a population, and equals another L-gate only when both exclude the same quadrant. The −1 must be read as this remainder and not from the signs of the components, since products of first-quadrant primes can lie anywhere in the plane: a CD14− CD16− quadrant has the step code −1829615469−2414358686i, with unit 1, whereas the L-gate excluding it is 1829615469+2414358686i, with unit −1. The remainder of −1 is valid only on a number built from exactly two axes; any other remainder, such as i or −i, is not part of the framework.

At the population level, however, the position of the −1 is lost. A population code multiplies the step codes of its gating path (§3.3), and a product keeps its factors but not their places: the −1 would become the unit of the whole number and would no longer show which two axes were excluded together. An L-gate is therefore never multiplied into a population code: it exists only as a step of the gating path, recorded in the fingerprint, and populations gated through it have no population code. Where every population needs its own code, the three remaining quadrants are gated separately, as CD45RA versus CCR7 and CD14 versus CD16 are in Supplementary Table S2.

### 2.3 Antibody clones recognizing multiple isoforms

When a clone recognizes several isoforms of the same protein, its identifier is the **product of the corresponding slot primes**. Clone `NCAM16.2`, which recognizes `P13591-1`, `-2` and `-3`, is therefore

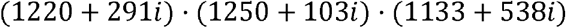

The slot primes are distinct, and the first-quadrant restriction of §2.1.2 fixes a single representative for each associate class. Factorizing the identifier therefore returns exactly the slots from which it was built, and so exactly the isoforms the clone recognizes. Divisibility carries the corresponding relation between reagents: the identifier of one clone divides that of another if and only if the first recognizes a subset of the isoforms recognized by the second, so comparing the reactivity of two clones is a single division.

### 2.4 Antibody clones recognizing carbohydrate epitopes

Some clones recognize a carbohydrate rather than a protein. HNK1, the CD57 clone of the panel in §3.2, binds a sulfated carbohydrate (8), and a carbohydrate has no entry in the reference dictionary, so none of the rules of §2.1 can give it an identifier. CD57 is not alone. The CD nomenclature of the Human Cell Differentiation Molecules (HCDM) organization, approved by the International Union of Immunological Societies and sanctioned by the World Health Organization, names the antigens recognized by clusters of validated antibodies (9). Most designations name a protein and reach the reference dictionary through its gene, but Table I of Engel et al. (9) classes nineteen as carbohydrates or glycosphingolipids: CD15, CD15s, CD15u, CD15su, CD17, CD57, CD60a, CD60b, CD60c, CD65, CD65s, CD75, CD75s, CD77, CD173, CD174, CD175, CD175s and CD176.

A carrier cannot stand in for the carbohydrate. The same carbohydrate sits on many glycoproteins and glycolipids, and the carriers remain on cells that lack it. HNK-1, the carbohydrate detected as CD57, is carried by NCAM, L1 and myelin-associated glycoprotein (10), yet CD56bright NK cells are CD57− while expressing NCAM, the protein detected as CD56 (11). Encoding CD57 by a carrier would make its negative state false. A carbohydrate therefore needs an identifier of its own.

The CD nomenclature itself supplies the list from which that identifier is drawn. A designation is a number, possibly followed by a suffix. A lower-case w before the number marks a provisional designation, and for carbohydrates a lower-case suffix marks a modification of the same carbohydrate sequence (9). The list used here is Table I of Engel et al. (9), 401 designations, updated with the eleventh HLDA workshop, which turned CDw198 into CD198 and added CD372 to CDw382 (12). It holds 412 designations under the numbers 1 to 382.

Both lists draw on the same primes, so the CD list must begin where the reference dictionary will not reach. A second constant is therefore fixed at publication: the reference dictionary is allotted 80,000 blocks, up to rank 5,120,000 (13544+759i). The reference release holds 585 entries created in 2016–2025, or 58.5 a year; at that rate the 59,569 blocks that follow the reference release last 1,018 years, far longer than this version of the standard is expected to serve. The primes after rank 5,120,000 form the **CD region** (Figure 3A). A third constant is its block depth, *c =* 1024: the designation with number *n* and suffix *s* owns slot *s* of block *n*, paired with the Gaussian prime of rank

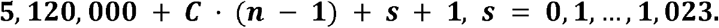

**Figure 3.**
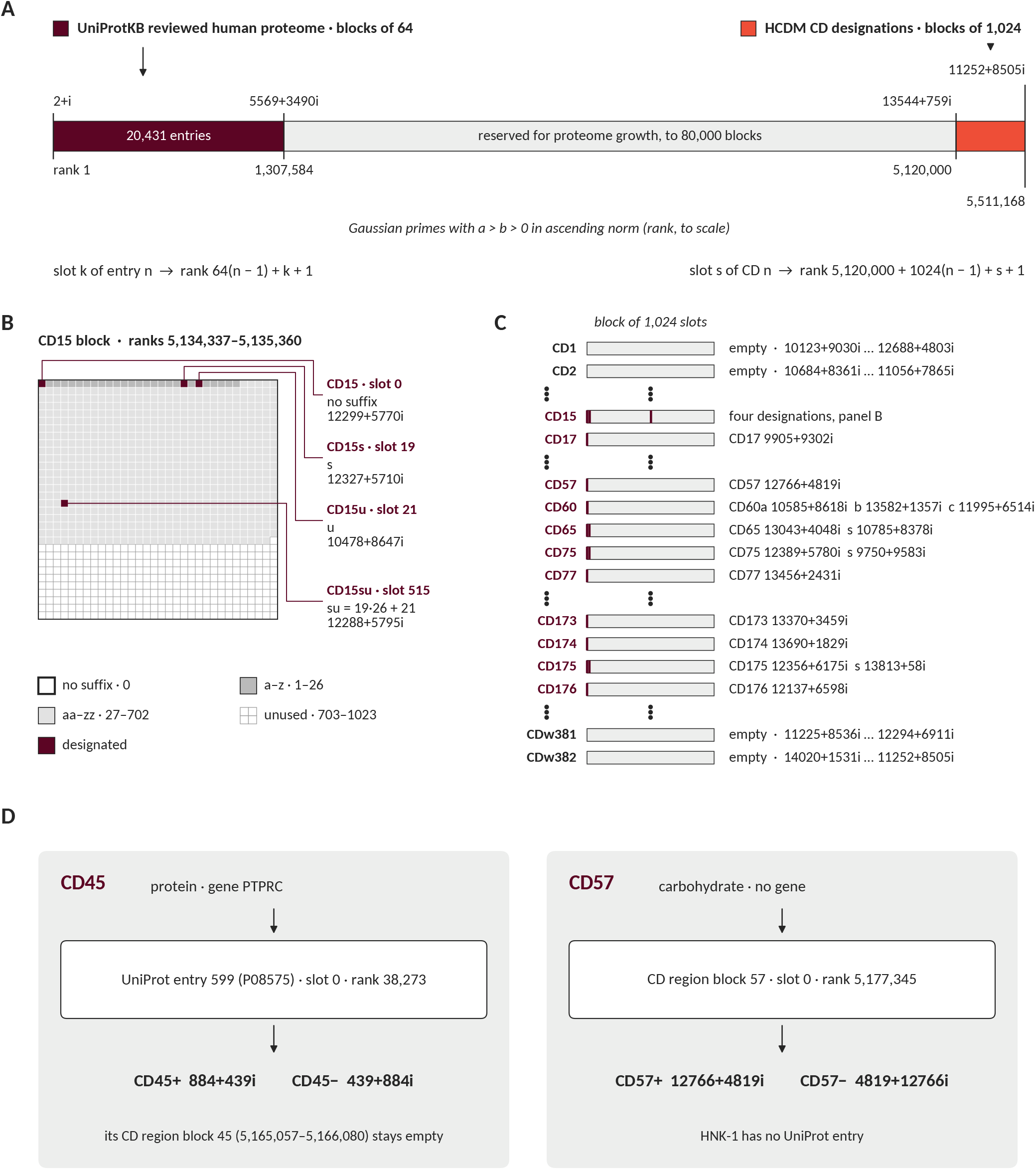
The CD region of the Gaussian prime dictionary. **(A)** The reference dictionary occupies ranks 1 to 1,307,584 (20,431 entries, blocks of 64) and is allotted the ranks up to 5,120,000 (80,000 blocks), about 1,000 years of growth at the 2016– 2025 rate of 58.5 entries a year. The CD region follows: each CD number owns a contiguous block of 1,024 primes, so slot *s* of CD *n* is the prime of rank 5,120,000 + 1,024(*n*−1) + *s* + 1. Ranks are drawn to scale. **(B)** Anatomy of one block, shown for CD15. Slot 0 holds the designation without suffix; a lower-case suffix is read as a spreadsheet column (a = 1 … z = 26, aa = 27 … zz = 702), so CD15s, CD15u and CD15su fall in slots 19, 21 and 515. **(C)** The CD region from CD1 to CDw382. Blocks of protein designations stay empty; only the 19 carbohydrate designations are assigned. **(D)** A designation with a gene is encoded in the reference dictionary, one without a gene in the CD region; negative states are mirrors, as in the reference dictionary. Primes are computed from the reference release (retrieved 23 March 2026), Table I of Engel et al. (9) and the designations of the eleventh HLDA workshop (12).

The slot index is read directly off the designation: *s* is 0 when there is no suffix, and otherwise the suffix read as a spreadsheet column, from a = 1 to z = 26 and from aa = 27 to zz = 702. Thus CD15 addresses slot 0 of block 15, CD15s slot 19, CD15u slot 21 and CD15su slot 19 · 26 + 21 = 515 (Figure 3B). As with isoform suffixes (§2.1.3), nothing intervenes between the designation and its slot. C = 1,024 is the smallest power of two that holds every suffix of up to two letters; the longest carbohydrate suffix, su, has two. Every number owns its block whether or not it carries a designation, so a new designation finds its address without consulting any list. Each designation is placed by the following rules.

Write each designation as CD*n* followed by its suffix *s*, ignoring a w before the number, and read its class in the HCDM list:

**(Carbohydrate)** If the class is carbohydrate or glycosphingolipid, the designation occupies slot *s* of block *n* and takes the Gaussian prime of that slot.

For example, CD57 occupies slot 0 of block 57, 12766+4819i, and its negative state is the mirror 4819+12766i (§2.2.1).

**(Protein)** If HCDM assigns the designation a gene, the protein is encoded in the reference dictionary by §2.1. Slot *s* of block *n* stays reserved and is never used, so no marker carries two identifiers. For example, CD45 is encoded by P08575, 884+439i, and block 45 of the CD region stays empty (Figure 3D).

**(Not cloned)** If HCDM lists the antigen as not cloned, the designation receives no identifier until a gene is assigned, and is then encoded under (Protein).

An address, once allocated, is permanent. A designation that HCDM later drops or reassigns keeps its slot as a tombstone, which is never reused, and confirming a provisional designation does not move it, since the w plays no part in the address.

Applied to the current list, these rules classify every designation without exception, and the 412 designations fall into three cases (Figure 3C).

#### Case 1

**names a carbohydrate** (19 designations in 11 blocks). The 19 identifiers run from CD15, 12299+5770i, to CD176, 12137+6598i; no component exceeds 13,813 and no norm exceeds 190,840,373.

#### Case 2

**names a protein** (389 designations, 7 of them splice variants such as CD45RA). A splice variant is encoded through the isoform slots of its gene (§2.1.5, §2.3). The 331 blocks that hold only protein designations stay empty.

#### Case 3

**names an antigen not cloned** (4 designations: CD139, CDw145, CD165 and CD245). Their blocks stay empty.

A further 36 numbers between 1 and 382 carry no current designation, and their blocks also stay empty. Block 382, the last, ends at rank 5,511,168 (11252+8505i; Figure 3A).

A clone clustered under a carbohydrate designation takes that designation’s identifier, so HNK1 is encoded by 12766+4819i. A clone that recognizes two designations takes their product, as for multiple isoforms (§2.3). A new designation takes its slot by the same rules and displaces nothing, so the CD region, like the reference dictionary, is append-only (§2.6).

The rules rest on one principle: **a carbohydrate is identified by its designation, not by the molecules that carry it**. HCDM decides what a designation names; the designation alone decides where it sits.

### 2.5 Vectorized representation and cryptographic hashing of hierarchical gating

To avoid the integer explosion from computing theoretical Gödel numbers, the method uses a lazy-evaluation approach, representing the gating strategy as a one-dimensional array rather than performing the final exponentiation. As a user defines a hierarchical gating strategy, the software computes the PPS identifier, a complex number, for each sequential gate, and the strategy is held in memory as an ordered array [PPS_1_, PPS_2_, …, PPSI⍰] of all gated cell populations. Because the Gödel sequence primes (2, 3, 5, 7, …) are implicit in the hierarchical steps, they are omitted to reduce overhead.

To generate the final immutable identifier for global database storage, the local array is transformed by deterministic cryptography. The array of complex numbers is first converted into a strictly formatted, universally standardized UTF-8 string; a three-step gating array, for example, is concatenated as *(a*_*1*_*+ b*_*1*_*i)(a*_*2*_*+ b*_*2*_*i)(a*_*3*_*+ b*_*3*_*i)*. As one number, an L-gate is one element of the gating array, and its string carries the signs of its components, so two paths that differ only in one step, the L-gate in one and the quadrant it excludes in the other, have different fingerprints. The string is then passed through SHA-256 (13), returning a fixed-length 64-character hexadecimal string. This collision-resistant hash serves as the unique universal identifier for the gating strategy, suitable for publication, sharing and relational database indexing.

### 2.6 Stability and growth

The identifier system is append-only. *A newly reviewed entry* carries a creation date later than every entry in the reference release and therefore sorts to the end. It takes index *n*+1, whose block of 64 primes is fixed by the same rule and belongs to no other entry. *A newly annotated isoform* of an existing protein takes the slot indexed by its UniProt suffix inside its parent’s own block. Neither event shifts a single existing assignment, so identifiers published under the reference release remain valid indefinitely and existing (active and deprecated) SHA-256 fingerprints remain verifiable.

## 3. Results

### 3.1 Objective generation of the universal UniProt dictionary

Pairing the reference release with the ordered prime list assigns an identifier to every protein and every isoform in the reviewed human proteome, with no human decision at any step. The procedure allocates 53,296 identifiers: 20,431 protein-level addresses and 32,865 isoform-level addresses. All are distinct, and every isoform description in the release, including all 329 descriptions carrying an accession prefix other than the citing entry, resolves under the single rule of §2.1.5. Spreading each identifier over the real and imaginary components keeps those components small (Figure 2). Across the 53,296 allocated identifiers no real or imaginary component exceeds 6,562 and no norm exceeds 43,191,689.

### 3.2 Assignment of a mass cytometry panel to Gaussian primes

We applied the rules of §2.1–2.4 to the 33 surface markers of the mass cytometry dataset of Scafidi et al. (14) that includes 30 antibodies of the Maxpar Direct Immune Profiling Assay (15) and three drop-in antibodies against CD33, TIGIT and CD279 (PD- 1). The assignment is given in Supplementary Table S1. All 33 markers were assigned. Twenty-nine clones take slot 0, the protein-level address (§2.1.4). Three clones are encoded by more than one prime: NCAM16.2 by the three NCAM1 isoforms it recognizes, 3G8 by FCGR3A and FCGR3B, since it binds both FcγRIIIa and FcγRIIIb (16), and HI100 by the four isoforms of exon A. One clone, HNK1, takes its identifier from the CD region (§2.4).

Three antibodies need comment:

#### CD45RA

The panel contains three anti-CD45 clones (Supplementary Figure S1). HI30 recognizes all CD45 isoforms and takes slot 0 (17). UCHL1 recognizes only CD45RO (18), which is `P08575-4`, and takes slot 4. HI100 belongs to the CD45RA cluster (17), and the CD45RA determinant requires the sequence encoded by exon A (19). Four isoforms retain exon A, `P08575-3`, `-5`, `-6` and `-8`, so HI100 is encoded as the product of the primes of slots 3, 5, 6 and 8 (§2.3), not by slot 8 alone, the isoform that UniProt names CD45RA.

#### TCRγδ

Clone B1 binds the γδ T-cell receptor, but its epitope has not been mapped (20). The δ chain has a single constant region, TRDC, and the γ chain has two, TRGC1 and TRGC2, either of which pairs with δ (21). We therefore encoded B1 by TRDC, the only chain present in every γδ T-cell receptor (Supplementary Figure S2). The identifier records the receptor that the clone binds, not its epitope, just as slot 0 records the protein for clones of undetermined isoform reactivity (§2.1.4). The clones against CD3 and HLA-DR were encoded in the same way, by CD3E and HLA-DRA.

#### CD57

CD57 is detected by HNK1, which recognizes a sulfated carbohydrate carried by several glycoproteins (8). A carbohydrate has no entry in the reference dictionary, so CD57 is encoded in the CD region (§2.4) by slot 0 of block 57: 12766+4819i for CD57+ and 4819+12766i for CD57−.

### 3.3 Assignment of unique identifiers to a cytometry gating strategy

We applied the rules of §2.2.1, §2.3, §2.4 and §2.5 to a gating strategy modified from Scafidi et al. (14), with markers encoded as in §3.2. Supplementary Table S2 shows the hierarchical gating strategy and the identifiers of every gate. The gating strategy comprises 264 marker gates, excluding the five technical gates on calibration beads, acquisition time, singlets and viability. Each gate receives a step code and a population code, and its gating path a fingerprint (Supplementary Table S2). The step code is the product of the markers used to define the gate. The population code is the product of the step codes along its gating path (gating hierarchy). The population code therefore equals the population code of its parent multiplied by its own step code. The ordered list of step codes along the path is the array of §2.5, and its SHA-256 hash is the fingerprint of the path (gating hierarchy). Figure 4 follows one gating path through its four steps (s_1_–s_4_): CD45+ CD66b− cells, CD3− CD19+ B cells, HLA-DR+ CD20+ B cells and, finally, IgD+ CD27+ unswitched memory B cells.

**Figure 4.**
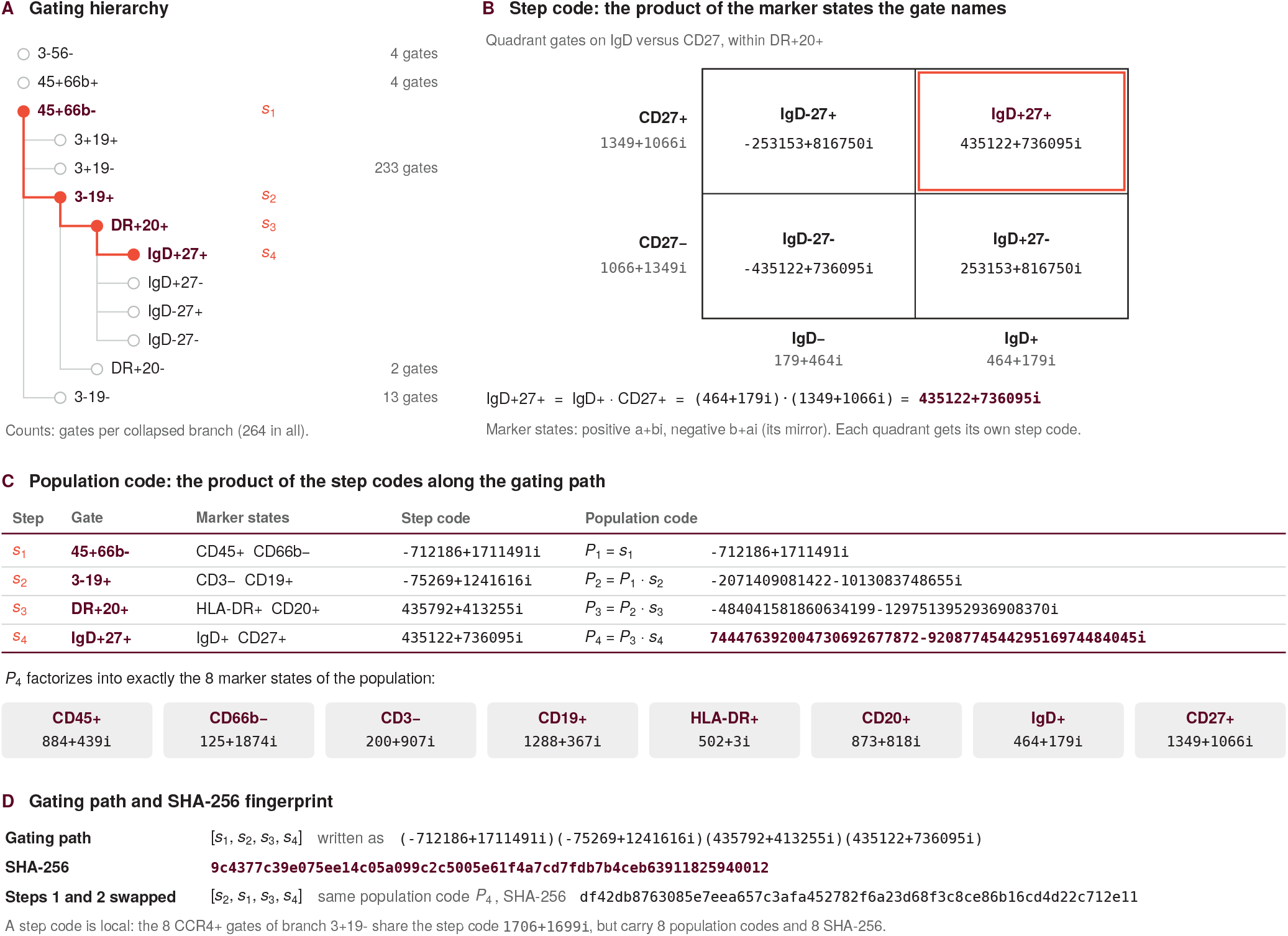
Identifiers of a cytometry gating strategy. **(A)** Gating hierarchy of the gating strategy modified from Scafidi et al. (14): 264 marker gates, with collapsed branches giving their number of gates. Technical gates are omitted. One gating path is highlighted, its steps labelled *s*_1_–*s*_4_. **(B)** The step code is the product of the marker states the gate names, each marker contributing its positive (a+bi) or negative (b+ai) state, so the four quadrant gates on IgD versus CD27 carry four step codes. **(C)** The population code is the product of the step codes along the gating path. Each population code is its parent’s multiplied by the step code, and factorization returns exactly the marker states of the population. **(D)** The ordered step codes are written as a string and hashed with SHA-256. Exchanging the first two steps leaves the population code unchanged but changes the fingerprint. A step code depends on the gate alone, so the eight CCR4+ gates share one, while their population codes and fingerprints differ. All gates are listed in Supplementary Table S2.

All 264 gates were encoded, with gating paths of one to seven steps. They carry 264 distinct population codes and 264 distinct fingerprints, but only 69 distinct step codes, because a step code depends on the gate alone. The eight CCR4+ gates of the T-cell branch, for example, share the step code 1706+1699i, while their population codes and fingerprints all differ. The largest population codes, those of the 128 chemokine- receptor subsets of CD4 and CD8 T cells, are products of 19 Gaussian primes. Every code was checked by factorization against the full dictionary, CD region included. Each of the 330 distinct codes returned exactly the identifiers and states from which it was built.

The population code and the fingerprint identify different things. The population code is a product, so it does not depend on the order of the steps, and factorizing it returns exactly the marker states that define the population. The fingerprint is computed on the ordered array, so it changes with the order. Exchanging the first two steps of the path in Figure 4 leaves the population code unchanged but changes the fingerprint. The population code thus identifies the cells, and the fingerprint identifies the gating path that reached them.

Two kinds of gates need comment:

#### Technical gates

The five gates on calibration beads, acquisition time, singlets and viability involve no protein and set no marker state. They carry no identifier and are not part of any gating path, which therefore starts at the first marker gate.

#### CD57

The eight CD57+ gates take CD57+ from the CD region (§2.4), 12766+4819i, as their step code. Like the CCR4+ gates, they share one step code, while their population codes and fingerprints differ.

## 4. Discussion

Building on the arithmetical foundations of the Prime Population System and Gödel numbering (1,2), this study introduces **Gaussian Prime Cytometry**, an identifier system in which every protein and isoform of the reference proteome is paired with a Gaussian prime. The two earlier layers keep their names and their scope, the PPS for cell populations and the UGI for gating strategies, but both are now built on Gaussian primes. Where the distinction matters we call them the Gaussian PPS and the Gaussian UGI, because their codes differ from those of the 2018 and 2025 definitions.

Because every identifier is a product of Gaussian primes, relationships between populations and reagents can be derived from simple arithmetical operations. The population code of every cell population descendant is divisible by that of its ancestor, and the quotient returns the marker states that separate them; more generally, one population code divides another whenever the first population is defined by a subset of the marker states of the second, whatever the gating path. The greatest common divisor of two population codes returns the marker states they share, and so their closest common phenotype. The same relation holds between reagents: the identifier of one clone divides that of another if and only if the first recognizes a subset of the isoforms recognized by the second (§2.3). Because *a+ bi* and *b+ ai* are distinct primes, these operations keep positive and negative states apart, while Conjugate Pair Symmetry still allows them to be paired. Each operation is exact and needs neither text parsing nor curated synonyms.

The price of the encoding is size. Spreading each identifier over two components keeps each component small (at most 6,562, against 229,937 for a dense assignment of the same 20,431 entries to rational primes), but the norms are larger. A negative state costs no more than a positive one, instead of the squared prime of the original system.

Population codes grow by about 10 to 12 bits per marker; most exceed 64 bits (247 of the 264 in Supplementary Table S2) and are therefore stored exactly, as text of the form a + bi or as arbitrary-precision integers. The gating path carries no such cost: its SHA- 256 fingerprint is always 64 hexadecimal characters, whatever the number of steps, whereas a Gödel integer grows with the values of the PPS numbers it uses as exponents.

Encoding a real gating strategy exposed two limitations. First, the five technical gates on calibration beads, acquisition time, singlets and viability set no marker state and carry no identifier, so every encoded path starts at the first marker gate. Population codes and fingerprints therefore describe the biological gating only: two analyses that share a fingerprint may still differ in how beads, unstable acquisition, doublets or dead cells were removed, and these gates must be reported separately. Second, step codes are read from the gate names, so they record the marker states the analyst declared, not where the gates were drawn. Two analyses that place a gate at different boundaries share its code, and a mislabelled gate passes its error to every code below it.

Gating-ML 2.0, the ISAC standard for exchanging gates between software tools (24), records what the codes leave out: the type and boundaries of each gate, its place in the gating sequence, and the compensation and scale of each axis, so that applying the file to the same data returns the same cells. What it does not record is the biology. Axes are named after instrument channels without any convention, and no field states which marker state a gate selects, so Gating-ML stays within the experimental domain (2). The two descriptions are therefore complementary. Written into the custom metadata of each Gating-ML gate, the step code, the population code and the fingerprint would link the experimental description to the biological one: two analyses that share a fingerprint could be compared gate by gate, and a gate whose axes are not the markers named in its step code could be flagged. This requires software that exports complete Gating-ML, including the scale definitions (24).

Matching fingerprints show that two gating paths declare the same marker states, on the same isoforms and in the same sequence, so datasets can be aligned by comparing hashes rather than by manually curating population names; a match marks candidate equivalent populations, which the gate definitions then confirm. This is possible because the dictionary is generated rather than curated: two lists, each ordered by a key intrinsic to it, pair by position alone, and the 20,431 protein-level and 32,865 isoform- level identifiers follow without a single act of expert judgement, so that every laboratory derives the same codes, as the FAIR principles require. The same holds true for the 412 designations of the CD nomenclature. A community registry pairing fingerprints with their arrays and readable names would make the hashes reversible.

It is worth being precise about what such a registry is and is not. The dictionary itself requires no registry: the pairing is determined by two public orderings and can be regenerated from the reference release by anyone, which is the point of §2.1. SHA-256 lookup is a convenience for reversing the hash of a gating strategy, not a dependency of the identifier scheme. By tethering the Gaussian primes to UniProt and to the CD nomenclature, the system eliminates the need for human curation or subjective consensus at the level where ambiguity arises.

## Supplementary material

**Supplementary Table S1. Assignment of 33 markers to Gaussian primes**. One row per identifier. Entry index *n*, slot *k* and rank 64(*n*−1)+*k*+1 follow §2.1; the positive state a+bi and the negative state b+ai follow §2.2.1. CD57, a carbohydrate, has no UniProt entry and takes slot 0 of block 57 of the CD region (§2.4): its row gives the CD number as *n* and the rank 5,120,000+1,024(*n*−1)+*k*+1. A clone listed on several rows is encoded as the product of those rows, in both states (§2.3). The isoform record gives the §2.1.5 case of each entry. Notes refer to the three antibodies discussed in §3.2. Identifiers are allocated against the reference release (UniProtKB/Swiss-Prot, *Homo sapiens*, retrieved 23 March 2026), and the CD57 identifier against the CD list of §2.4.

**Supplementary Figure S1. Reactivity of the anti-CD45 clones HI30, UCHL1 and HI100 across the eight PTPRC isoforms**. Rows list the isoforms of UniProtKB P08575 in slot order; the slot is the integer suffix of the UniProt isoform identifier. Isoform names and variable-exon composition are taken from the UniProt entry. Filled squares mark the variable exons A, B and C (exons 4–6) retained in each isoform; exon A (orange) carries the CD45RA determinant. The Gaussian prime column gives the identifier allocated to each slot. Dots mark the isoforms bound by each clone. HI30 recognizes a determinant common to all isoforms (17) and takes slot 0. UCHL1 binds only CD45RO, the 180 kDa isoform lacking all variable exons (18), and takes slot 4. HI100 belongs to the CD45RA cluster (17), whose determinant requires exon-A- encoded sequence (19); it is encoded as the product of the identifiers of slots 3, 5, 6 and 8.

**Supplementary Figure S2. Encoding of clones against receptor complexes**. Rows list the forms of each receptor and columns its chains, with the UniProt accession and slot-0 Gaussian prime of each chain in the reference release. Filled squares mark the chains present in each form. Each clone is encoded by the chain present in every form of its receptor (orange), regardless of which chain carries its epitope. The CD3 antigen comprises the CD3εγ and CD3εδ dimers (22); HLA-DR pairs its non-polymorphic α chain with a β chain encoded by one of the DRB genes (23); the γδ T-cell receptor pairs its δ constant region with either γ constant region (21). UCHT1, LN3 and B1 are therefore encoded by CD3E, HLA-DRA and TRDC. The white dot marks the only epitope chain documented in the literature, CD3ε for UCHT1 (22).

**Supplementary Table S2. Identifiers of the gates of the gating strategy**. One row per marker gate (264) of the gating strategy modified from Scafidi et al. (14), in the order of the FlowJo export; the five technical gates are omitted (§3.3). The step code is the product of the marker states named by the gate. The population code is the product of the step codes along the gating path, given with the number of its Gaussian prime factors. Steps 1 to 7 give the step codes of the gating path in order. SHA-256 gives the fingerprint of the path, computed on the string (a_1_+b_1_i)(a_2_+b_2_i)… (§2.5). Marker states follow Supplementary Table S1. CD57 takes its identifier from the CD region (§2.4).

Further sheets list the positive and negative states of the 30 markers named in the strategy, the verification checks and the rules applied.

## Databases

### PPS_Database1

PPS Database 1 includes the 20,431 entries together with the primes assigned to the slots. It does not contain the isoform assignment but contains the complete Gaussian prime list including the not assigned primes. https://public.tableau.com/views/PPS_Database1/ReferenceReleasePairing?:language=en-US&:sid=&:redirect=auth&:display_count=n&:origin=viz_share_link

### PPS_Database2

PPS Database 2 includes the complete UniProt assignment. Each tab shows some examples of data as they are described in §2.1.4 and §2.1.5. https://public.tableau.com/views/PPS_Database2_V2/Entries?:language=en-US&:sid=&:redirect=auth&:display_count=n&:origin=viz_share_link

PPS_CD_Database2. PPS CD Database 2 includes the complete CD nomenclature assignment. The first tab shows the complete list of CD markers as well as some visualizations of the 19 carbohydrates with a Gaussian prime assigned. https://public.tableau.com/app/profile/lihpublicdata/viz/PPS_CD_Database2/CDlist

## Declaration of generative AI use

In preparing this work, the authors used Claude Opus 5.5 (Anthropic) to write the scripts that build and check the Gaussian prime dictionary, to produce all figures, and to assist with drafting and editing the text. The operations performed by these scripts are arithmetic, and their output was spot-checked by the authors. The authors verified the dictionary and gate identifiers by spot checks against the rules of §2.1–2.5. All figures were rendered by these scripts from computed data rather than generated as images by the AI tool. The authors reviewed and edited all content and take full responsibility for the published article.

## Literature Cited

1. Cosma A. Universal cell type identifier based on number theory. Cytometry A 2018;93:649–652.

2. Cosma A. Unique gating strategy identifier based on the Prime Population System and Gödel numbers. Front Immunol 2025;16:1481412.

3. Weissman MH. An Illustrated Theory of Numbers. Providence, RI: American Mathematical Society; 2017.

4. UniProt Consortium. UniProt: The Universal Protein Knowledgebase in 2025. Nucleic Acids Res 2025;53:D609–D617.

5. Hardy GH, Wright EM. An Introduction to the Theory of Numbers. 6th ed. Revised by Heath-Brown DR, Silverman JH. Oxford, UK: Oxford University Press; 2008.

6. Sallusto F, Lenig D, Förster R, Lipp M, Lanzavecchia A. Two subsets of memory T lymphocytes with distinct homing potentials and effector functions. Nature 1999;401:708–712.

7. Ziegler-Heitbrock L, Ancuta P, Crowe S, Dalod M, Grau V, Hart DN, Leenen PJ, Liu YJ, MacPherson G, Randolph GJ, et al. Nomenclature of monocytes and dendritic cells in blood. Blood 2010;116:e74–e80.

8. Kizuka Y, Oka S. Regulated expression and neural functions of human natural killer-1 (HNK-1) carbohydrate. Cell Mol Life Sci 2012;69:4135–4147.

9. Engel P, Boumsell L, Balderas R, Bensussan A, Gattei V, Horejsi V, Jin BQ, Malavasi F, Mortari F, Schwartz-Albiez R, et al. CD nomenclature 2015: Human leukocyte differentiation antigen workshops as a driving force in immunology. J Immunol 2015;195:4555–4563.

10. Kruse J, Mailhammer R, Wernecke H, Faissner A, Sommer I, Goridis C, Schachner M. Neural cell adhesion molecules and myelin-associated glycoprotein share a common carbohydrate moiety recognized by monoclonal antibodies L2 and HNK-1. Nature 1984;311:153–155.

11. Nielsen CM, White MJ, Goodier MR, Riley EM. Functional significance of CD57 expression on human NK cells and relevance to disease. Front Immunol 2013;4:422.

12. Fernández-Calles J, Kužílková D, Hedin F, Bakardjieva-Mihaylova V, Škvárová Kramarzová K, van Zelm MC, Cosma A, Kalina T, Engel P. CD molecules nomenclature 2025: Antibody validation and expression profiling of immune system G protein-coupled receptors. Eur J Immunol 2025;55:e70099.

13. National Institute of Standards and Technology. Secure Hash Standard (SHS). FIPS PUB 180-4. Gaithersburg, MD: National Institute of Standards and Technology; 2015.

14. Scafidi A, Kaoma T, Cerella C, Campus E, Grzyb K, Nosirov B, Lind-Holm Mogensen F, Klein E, Da Costa R, Skupin A, et al. Circulating immune profiling reveals impaired monocyte states and trajectories driving immunosuppression in glioblastoma. J Neuroinflammation 2026;23:306.

15. Standard BioTools. Maxpar Direct Immune Profiling Assay (MDIPA): Human whole blood and PBMC immunophenotyping panel, 30 markers. South San Francisco, CA: Standard BioTools.

16. Vely F, Gruel N, Moncuit J, Cochet O, Rouard H, Dare S, Galon J, Sautes C, Fridman WH, Teillaud JL. A new set of monoclonal antibodies against human FcγRII (CD32) and FcγRIII (CD16): Characterization and use in various assays. Hybridoma 1997;16:519–528.

17. Schwinzer R. Cluster report: CD45/CD45R. In: Knapp W, Dörken B, Gilks WR, et al., editors. Leucocyte Typing IV: White Cell Differentiation Antigens. Oxford, UK: Oxford University Press; 1989. pp 628–634.

18. Terry LA, Brown MH, Beverley PC. The monoclonal antibody, UCHL1, recognizes a 180,000 MW component of the human leucocyte-common antigen, CD45. Immunology 1988;64:331–336.

19. Streuli M, Morimoto C, Schrieber M, Schlossman SF, Saito H. Characterization of CD45 and CD45R monoclonal antibodies using transfected mouse cell lines that express individual human leukocyte common antigens. J Immunol 1988;141:3910–3914.

20. Dutta I, Postovit LM, Siegers GM. Apoptosis induced via gamma delta T cell antigen receptor “blocking” antibodies: A cautionary tale. Front Immunol 2017;8:776.

21. Krangel MS, Band H, Hata S, McLean J, Brenner MB. Structurally divergent human T cell receptor gamma proteins encoded by distinct C gamma genes. Science 1987;237:64–67.

22. Arnett KL, Harrison SC, Wiley DC. Crystal structure of a human CD3-ε/δ dimer in complex with a UCHT1 single-chain antibody fragment. Proc Natl Acad Sci U S A 2004;101:16268–16273.

23. Andersson G. Evolution of the human HLA-DR region. Front Biosci 1998;3:d739–d745.

24. Spidlen J, Moore W, ISAC Data Standards Task Force, Brinkman RR. ISAC’s Gating-ML 2.0 data exchange standard for gating description. Cytometry A 2015;87:683–687.

