## Supplementary Figure S1 for "A Multidimensional Arithmetical Ontology for High-Parameter Cytometry: Gaussian Primes and Cryptographic Hashing for Cell Population Standardization"

| CD45 isoform | UniProt ID | Slot | Variable exons | | | Gaussian prime | Clone reactivity | | |
|---|---|---|---|---|---|---|---|---|---|
| | | | A | B | C | | HI30 | UCHL1 | HI100 |
| **CD45RABC** | P08575-3 | 3 | ■ | ■ | ■ | 932+325i | ● | | ● |
| **CD45RO** | P08575-4 | 4 | □ | □ | □ | 905+394i | ● | ● | |
| **CD45RAB** | P08575-5 | 5 | ■ | ■ | □ | 987+10i | ● | | ● |
| **CD45RAC** | P08575-6 | 6 | ■ | □ | ■ | 968+193i | ● | | ● |
| **CD45RBC** | P08575-7 | 7 | □ | ■ | ■ | 967+198i | ● | | |
| **CD45RA** | P08575-8 | 8 | ■ | □ | □ | 979+126i | ● | | ● |
| **CD45RB** | P08575-9 | 9 | □ | ■ | □ | 948+275i | ● | | |
| **CD45RC** | P08575-10 | 10 | □ | □ | ■ | 876+455i | ● | | |

**HI30**
all eight isoforms
**slot 0 (protein level)**

**UCHL1**
CD45RO only
**slot 4**

**HI100**
four exon-A isoforms
**slots 3, 5, 6, 8**

■ exon A retained (CD45RA determinant) ■ exon B or C retained □ exon spliced out ● ● ● clone binds
