## Supplementary Figure S2 for "A Multidimensional Arithmetical Ontology for High-Parameter Cytometry: Gaussian Primes and Cryptographic Hashing for Cell Population Standardization"

| **CD3 antigen** clone UCHT1 | **CD3E** P07766 907+200i | **CD3G** P09693 853+670i | **CD3D** P04234 667+242i |
|---|---|---|---|
| CD3εγ dimer | chain present in every form, documented epitope chain | chain of this form only | chain absent |
| CD3εδ dimer | chain present in every form, documented epitope chain | chain absent | chain of this form only |

| **HLA-DR** clone LN3 | **HLA-DRA** P01903 502+3i | **HLA-DRB1** P01911 503+60i | **HLA-DRB3** P79483 3163+2822i | **HLA-DRB4** P13762 949+844i | **HLA-DRB5** Q30154 4589+4100i |
|---|---|---|---|---|---|
| α with DRB1 | chain present in every form | chain of this form only | chain absent | chain absent | chain absent |
| α with DRB3 | chain present in every form | chain absent | chain of this form only | chain absent | chain absent |
| α with DRB4 | chain present in every form | chain absent | chain absent | chain of this form only | chain absent |
| α with DRB5 | chain present in every form | chain absent | chain absent | chain absent | chain of this form only |

| **γδ T-cell receptor** clone B1 | **TRDC** B7Z8K6 5233+3692i | **TRGC1** P0CF51 6010+2213i | **TRGC2** P03986 658+83i |
|---|---|---|---|
| Cγ1 form | chain present in every form | chain of this form only | chain absent |
| Cγ2 form | chain present in every form | chain absent | chain of this form only |

**UCHT1**
CD3: CD3E in every form
**identifier P07766, slot 0**
epitope chain: CD3ε (documented)

**LN3**
HLA-DR: HLA-DRA in every form
**identifier P01903, slot 0**
epitope chain: not established

**B1**
γδ TCR: TRDC in every form
**identifier B7Z8K6, slot 0**
epitope chain: not documented

Legend: chain present in every form · chain of this form only · chain absent · documented epitope chain
